# Identification of a functional small non-coding RNA encoded by African swine fever virus

**DOI:** 10.1101/865147

**Authors:** Laura E. M. Dunn, Alasdair Ivens, Christopher L. Netherton, David A. G. Chapman, Philippa M. Beard

## Abstract

*African swine fever virus* (ASFV) causes a lethal haemorrhagic disease of domestic pigs, to which there is no vaccine available. ASFV has a large, double-stranded DNA genome that encodes over 150 proteins. Replication takes place in the cytoplasm of the cell and involves complex interactions with host cellular components including small non-coding RNAs (sncRNAs). A number of DNA viruses are known to manipulate sncRNA either by encoding their own or disrupting host sncRNA. In order to investigate the interplay between ASFV and sncRNAs, study of host and viral small RNAs extracted from ASFV-infected primary porcine macrophages (PAMs) was undertaken. We discovered that ASFV infection had only a modest effect on host miRNAs, with only 6 miRNAs differentially expressed during infection. The data also revealed 3 potential novel small RNAs encoded by ASFV, ASFVsRNA1-3. Further investigation of ASFVsRNA2 detected it in lymphoid tissue from pigs with ASF. Overexpression of ASFVsRNA2 led to up to a 1 log reduction in ASFV growth indicating that ASFV utilises a virally-encoded small RNA to disrupt its own replication. This study describes the modest impact of ASFV on host sncRNAs and the identification of a functional ASFV-encoded sncRNA.

**Importance:** African swine fever (ASF) poses a major threat to pig populations and food security worldwide. The disease is endemic in Africa and Eastern Europe and rapidly emerging into Asia where it has led to the deaths of millions of pigs in the past 12 months. The development of safe and effective vaccines to protect pigs against ASF has been hindered by lack of understanding of the complex interactions between ASFV and the host cell. We focused our work on characterising the interactions between ASFV and sncRNAs. We found only modest changes to host sncRNA abundance after ASFV infection, and discovered a functional ASFV-encoded sncRNA. The knowledge from this study can be exploited to develop more effective ASFV vaccines that take advantage of the sncRNA system.

## Introduction

African swine fever (ASF) is a highly pathogenic viral disease of swine. Virulent strains cause acute haemorrhagic fever in domestic pigs with mortality rates up to 100% (1). There is currently no effective vaccine or treatment (2). ASF is caused by African swine fever virus (ASFV), the only member of the Asfarviridae family. The virus sits within the nucleocytoplasmic large DNA virus (NCLDV) superfamily which also includes the Poxviridae. NCLDVs have large, double-stranded DNA genomes and replicate in the cytoplasm of infected cells (3). ASFV replicates predominantly in the cytoplasm of cells of the monocyte/macrophage lineage. ASFV evades the anti-viral defences in these cells by modulation of a number of host-cell pathways including type I IFN induction (4), apoptosis (5), host-cell protein synthesis (6), and the NF-κB and NFAT signalling pathways (7). Descriptions of these host-cell interactions are reviewed in detail in (8)

Small non-coding RNAs (sncRNAs) are classes of small RNA (<200nt) which are involved in the regulation of gene expression and genome stability, predominantly through RNA interference (RNAi) mechanisms. Eukaryotic cells produce multiple classes of sncRNA, including microRNAs (miRNAs), PIWI-interacting RNAs (piRNAs) and endogenous small interfering RNAs (siRNAs) (9). These sncRNAs are involved in many biological processes including apoptosis, differentiation, stress response and immune activation (10). It is therefore unsurprising that viruses manipulate and exploit sncRNAs for their own benefit. Virus-encoded miRNAs have been identified in a number of DNA virus families including Herpesviridae, Polyomaviridae, Iridoviridae, Ascoviridae, Baculoviridae and the Adenoviridae (11). These miRNAs play a variety of roles including cell proliferation regulation, control of apoptosis and modulation of host immunity (12). For example the Kaposi’s sarcoma-associated herpesvirus (KSHV) encoded miRNA, miR-K1, regulates the switch between lytic and latent viral replication by control of NF-κB expression via targeting of the host IκBα transcript (13). Other classes of viral ncRNAs have also been identified (reviewed in (14)). An interesting example is the Herpes simplex virus type 1 (HSV-1) encoded non-miRNA small RNAs (LAT sRNA1 and sRNA1) that regulate productive infection and inhibit apoptosis (15).

DNA viruses have also been shown to manipulate host sncRNAs by targeting specific host miRNAs as in the case of murine cytomegalovirus (MCMV) infection. MCMV induces degradation of cellular miR-27a and miR-27b which are important for MCMV replication *in vivo* (16). A more non-specific and global effect is wrought by Vaccinia virus (VACV), the prototypic poxvirus and NCLDV member, which induces widespread disruption of host miRNAs by a process of 3’ polyadenylation and decay (17, 18). As RNAi is the major antiviral pathway in invertebrates, a number of arthropod-borne (arboviruses) are known to manipulate sncRNA during replication to evade this immune response. Interestingly, ASFV is currently the only known DNA arbovirus, and replicates in the soft tick vector of the *Ornithodoros* spp., which have a functional RNAi system (19). Overall, it is apparent that manipulation of sncRNA systems is a common feature of viruses in order to further their survival, replication and pathogenesis. We therefore sought to investigate the interaction between ASFV and sncRNAs.

The effect of ASFV infection on host miRNAs has been investigated *in vivo* by comparing miRNA expression in pigs infected with a virulent strain to those infected with an attenuated strain (20). These authors identified 12 miRNAs that were differentially expressed. In addition a further study looked *in vivo* at the potential for ASFV to encode its own miRNAs and concluded that ASFV does not express miRNAs *in vivo* (21). In our study, we investigated the effect of ASFV infection on sncRNA in primary porcine alveolar macrophages *in vitro*. We found that virulent ASFV infection of primary porcine macrophages had only a small impact on host miRNAs with only 6 out of 178 identified porcine miRNAs differentially expressed over a 16 h time period. Interestingly, we discovered an ASFV-encoded sncRNA that, when overexpressed, led to a significant reduction in ASFV replication.

## Results

### ASFV infection does not induce polyadenylation or decay of cellular miRNAs

VACV, the prototypic poxvirus and NCLDV member, has been shown to induce widespread disruption of cellular miRNAs via a process of 3’ polyadenylation and decay (18) (17). In order to investigate if ASFV shares the ability of VACV to induce miRNA polyadenylation and decay porcine alveolar macrophages (PAMs) were infected with the pathogenic ASFV Benin 97/1 strain and Vero cells were infected with the Vero cell adapted ASFV strain, Ba71v. In parallel, PAMs and Vero cells were infected with VACV WR, with the samples collected and processed for northern blotting as described. The miRNA miR-27b-3p was selected as a probe for this experiment since it is extensively polyadenylated and degraded in VACV-infected cells (18). As expected, in VACV-infected Veros at 0 hpi mature miR-27b-3p was seen as a lower band (Fig 1a arrow) but by 6 hpi this mature form was almost undetectable and there was a higher molecular weight “smear” present in the lane, consistent with polyadenylation of the miRNA (Fig 1a asterisk). The higher molecular weight band above the smear in all lanes represents the precursor miRNA. By 16 hpi there was near complete decay of all forms of miR-27b-3p. In comparison, mature miR-27b-3p was present at all time points in ASFV-infected Veros, with no evidence of polyadenylation or decay. In VACV-infected PAMs there was visible polyadenylation of miR-27b-3p but an absence of decay. This is likely explained by VACV being unable to undergo a complete replication cycle in PAMs (data not shown). There was no modification or reduction detected in the amount of miR-27b-3p in ASFV infected PAMs. Levels of miR-27b-3p expression was quantified and normalised to 5s (Fig 1b). This highlighted the near 100-fold reduction in miR-27b-3p expression in VACV infected Veros whereas in ASFV-infected Veros and PAMs no reduction was detected.

**Figure 1:**
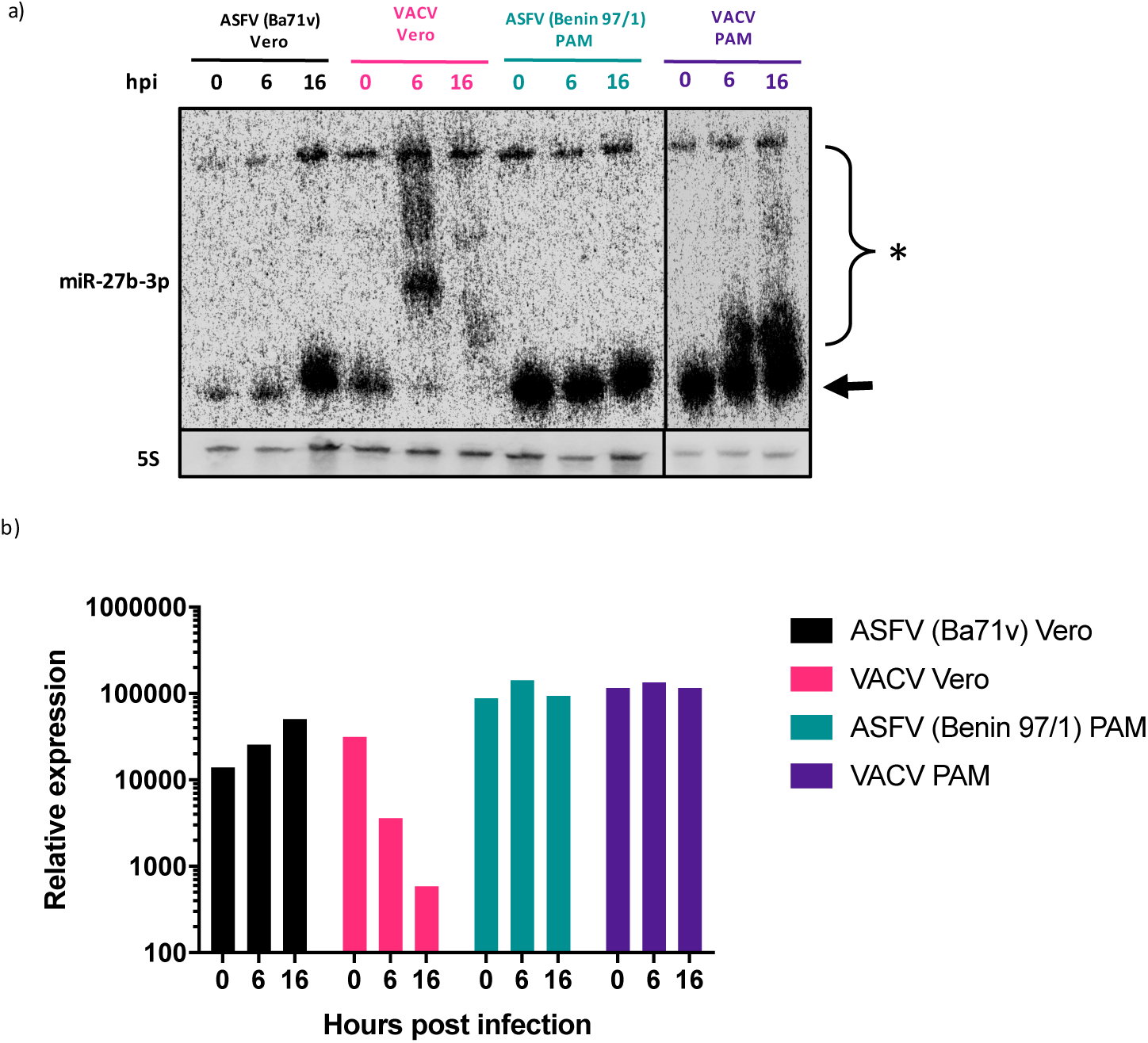
Infection with ASFV does not induce miRNA polyadenylation or decay. **a)** Northern blot of RNA extracted from Vero cells and porcine alveolar macrophages (PAMs) that were infected with either ASFV or VACV. The blot was probed for miR-27b-3p and 5s and imaged on a Phosphorimager. Arrow: mature miRNA, asterisk: polyadenylated miRNA **b)** Phosphorimager quantification of miR-27b-3p expression, normalised to 5S rRNA. The image and quantification is representative of 2 biological repeats.

In order to look comprehensively at the effect of ASFV infection on sncRNA, we utilised small RNA sequencing. RNA was extracted and sequenced from three biological repeats of either mock or ASFV-infected PAMs at 0, 6 and 16 hpi. Sequences aligning to 247 different mature miRNAs were obtained, these were then filtered to only include sequences with an average ≥ 5 reads as these gave higher quality boxplots, reducing the total number of miRNAs identified to 178. These accounted for, on average, 73% of total trimmed small RNA reads. Data from a previous study investigating the effect of VACV infection on host miRNAs was used as a comparison (18). In ASFV-infected cells there was no variation in the miRNA proportion of the total small RNA reads at both early and late time points compared to uninfected cells (Fig 2a). This was in contrast to VACV-infected cells, which had a 30% reduction in miRNA reads at an early time point (6 hpi) and over 50% reduction at late times (24 hpi) (Fig 2b) (18). To assess the extent of ASFV-induced 3’ modification of miRNAs, all trimmed sequences were analysed for non-templated nucleotide (nt) 3’ additions after nt position 19. No difference was detected between mock and ASFV infected samples at both 6 and 16 hpi, with the proportion of reads containing miRNAs with 3’ mismatches remaining at approximately 17% (Fig 2c). VACV infection led to a significant increase in the proportion of 3’ modified miRNA reads, which increased from 10% in mock to 25% in infected samples at 6 hpi (Fig 2d) (18). At 24 hpi, the difference was less substantial and increased from 10% in mock to 15% in infected samples, though was still statistically significant. (Fig 2d). In a final analysis of 3’ miRNA modification during ASFV infection, the extent of 3’ polyadenylation was examined by calculation of the proportion of miRNA reads which contained 3 or more non-templated 3’ adenosine residues beyond nt position 19. No difference was detected between mock and ASFV infected samples at either 6 or 16 hpi (Fig 2e). The results from northern blotting and small RNA sequencing revealed that ASFV does not share the ability of poxviruses to induce cellular miRNA polyadenylation and decay.

**Figure 2:**
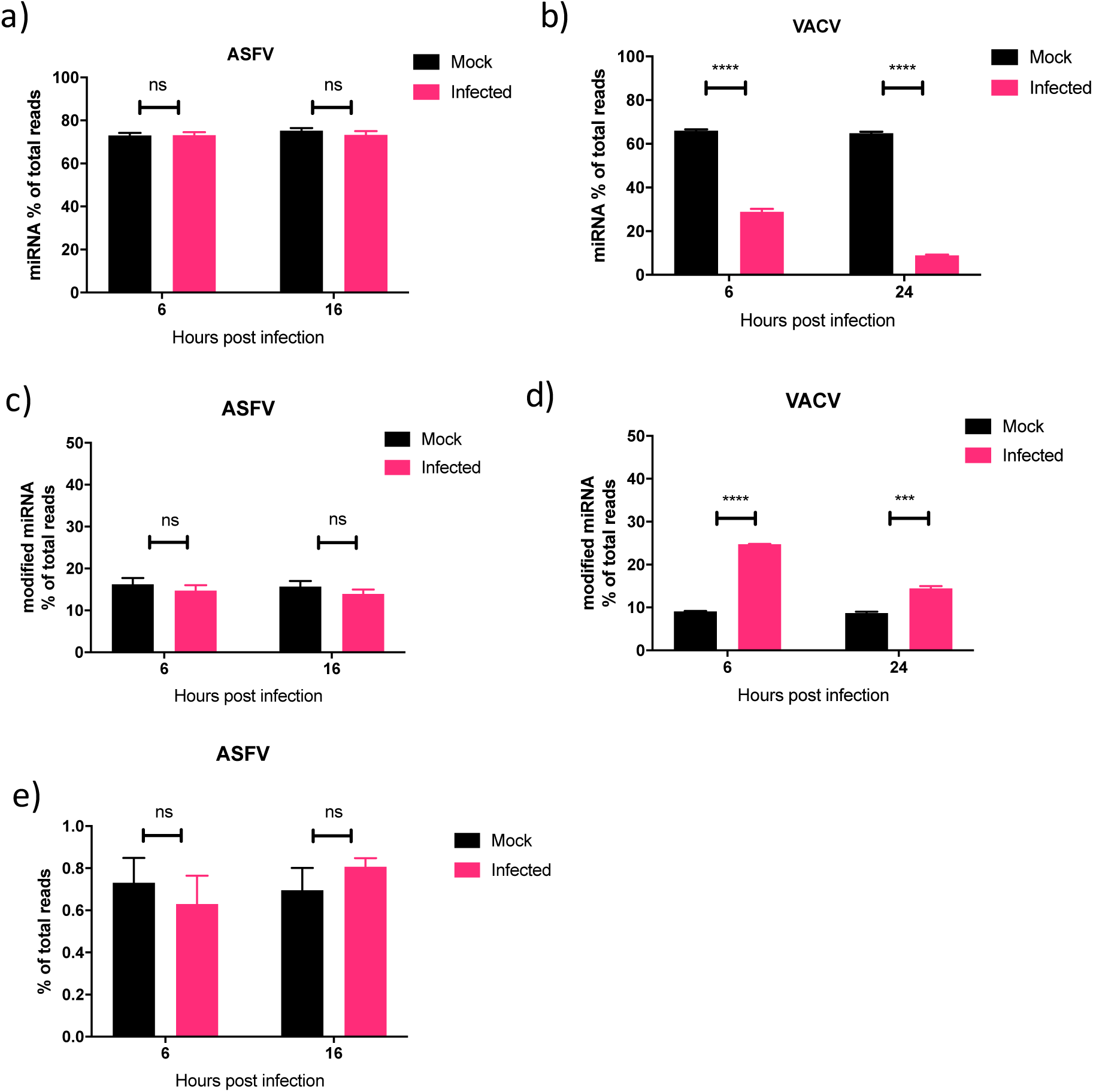
ASFV does not induce widespread polyadenylation or decay of cellular miRNAs. The proportion of sequencing reads that mapped to known host miRNAs a) PAMs infected or mock-infected with ASFV and in b) HeLa cells infected or mock infected with VACV at early and late time points. The proportion of 3’ modified miRNA, which contained at least 1 non-templated nucleotide after position 19 as a proportion total sequencing reads in c) ASFV mock and infected PAMs and d) VACV mock and infected HeLas. e) Proportion of miRNA containing 3 or more 3’ non-templated adenosine residues after nt position 19 in ASFV mock and infected PAMs. Data represent mean of 3 biological represents, error bars represent SEM. Data was statistically analysed using Student T-test: ns,p> 0.05, ***,p<0.001, ****,p<0.0001. Data from b) and d) is taken from supporting information from (18) and represents mean of 3 biological replicates.

### ASFV infection induces rapid changes in abundance of a select number of miRNAs

To determine if ASFV infection induces differential abundance of specific miRNAs we used the small RNA sequencing data to analyse changes in expression of individual miRNAs during infection, relative to mock infected cells. The filtered data (miRNA ≥5 reads, 178 miRNAs in total) was used for this analysis. MiRNAs were analysed for differential expression at 0, 6 and 16 hpi, relative to mock, and displayed on volcano plots (Fig 3a-c) for each time point. The majority of cellular miRNAs were not differentially expressed in response to ASFV infection. At both 6 and 16 hpi only one miRNA was significantly differentially expressed (Fig 3b, c). Expression of miR-10b increased 3.89 Log2 fold at 6 hpi and miR-27b-5p expression decreased 4.29 Log2 fold at 16 hpi. Interestingly, the time point with the most changes in miRNA expression was 0 h (Fig 3a). The 0 h samples were collected after the virus had been incubated on the cells for 1h at 37°C. Four miRNAs were significantly upregulated at this early time point: miR-10b, miR-486-1, miR-144 and miR-199a. Overall, the sequencing data indicated that ASFV infection does not have a widespread impact on host miRNA expression, but does lead to rapid changes in the abundance of a small number of miRNAs.

**Figure 3:**
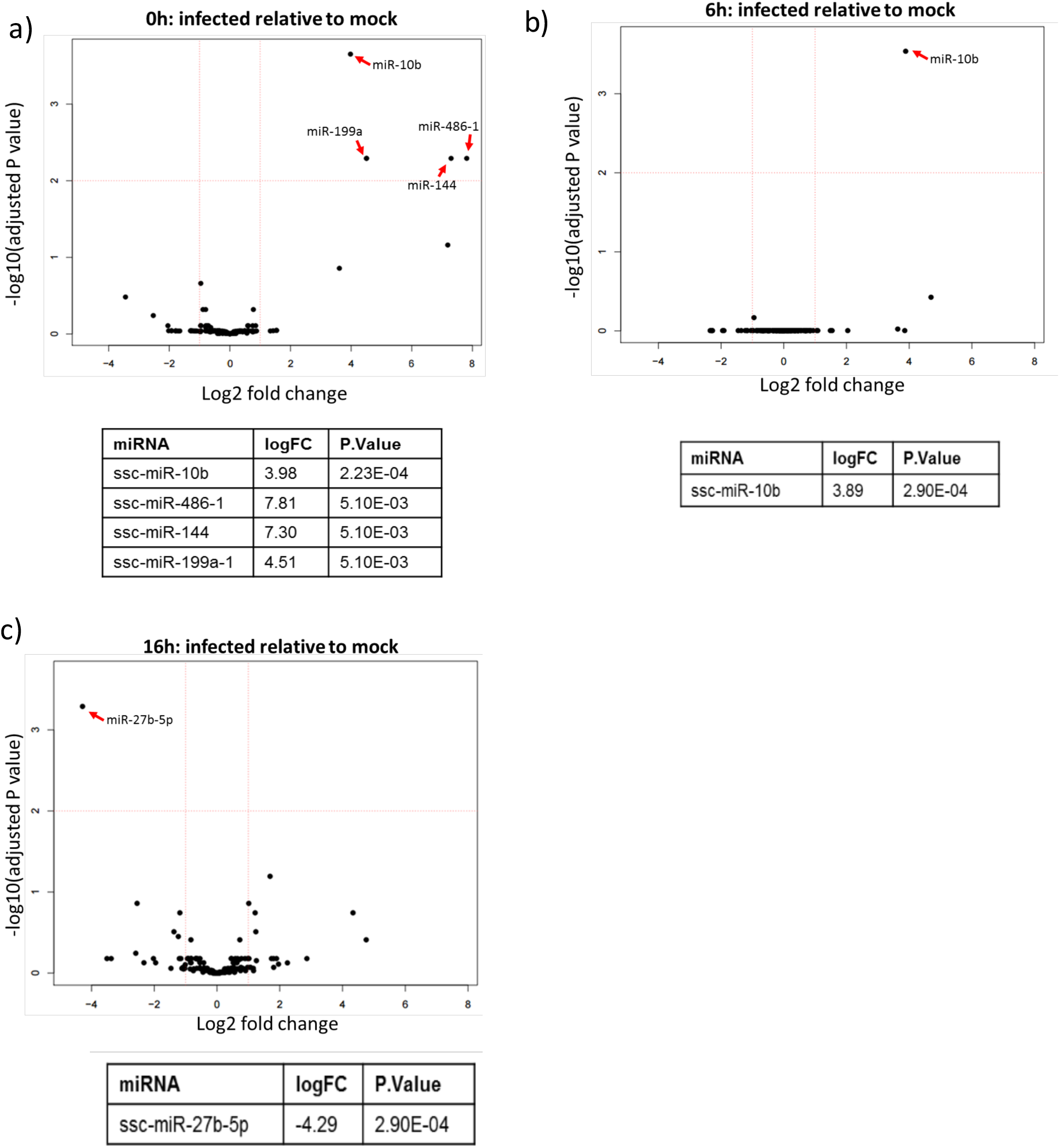
ASFV infection induces rapid changes in the abundance of a select number of miRNAs in PAMs. Volcano plots showing the differential expression of host miRNAs in ASFV infected cells relative to mock infected at 0 hpi (a), 6 hpi (b) and 16 hpi (c). The miRNAs that were differentially expressed with a significant adjusted P-value (≤0.05) are detailed in the table below each volcano plot.

In order to validate the miRNA expression changes identified from the sequencing analysis RT-qPCR was used to measure the abundance of individual miRNAs. Three more biological repeats of ASFV infections were repeated in PAMs (taken from 3 different pigs), RNA extracted at 0, 6 and 16 hpi and miRNA expression examined by RT-qPCR. To confirm that the miRNA expression changes were not the result of unspecific stimulation of macrophages due to cell debris in the virus preparation, we also tested miRNA expression by RT-qPCR after the addition of a mock virus preparation. In infected PAMs, RT-qPCR validated the upregulation of miR-10b at 0h to comparable levels detected in the sequencing (Fig 4a) however the upregulation at 6 hpi was not detected. Validation of miR-144 expression followed a similar pattern to the sequencing, though the upregulation was not as substantial (Fig 4b) with a log2 fold change of only 2.2 compared to 7.81 found by sequencing. RT-qPCR validation of miR-27b-5p expression showed a trend for downregulation but, again, this was not as substantial as detected by sequencing with only log2 fold change of −1, compared to −4 in the sequencing data at 16 hpi (Fig 4c). RT-qPCR was unable to validate miR-486-1 upregulation (Fig 4d). We were also unable to validate expression of miR-199a-1 due to its low expression in PAMs. With an average of only 17 miR-199a-1 reads per sample, a standard curve could not be generated. The mock virus preparation did not lead to the dysregulation of any of the tested miRNAs (Fig 4a-d).

**Figure 4:**
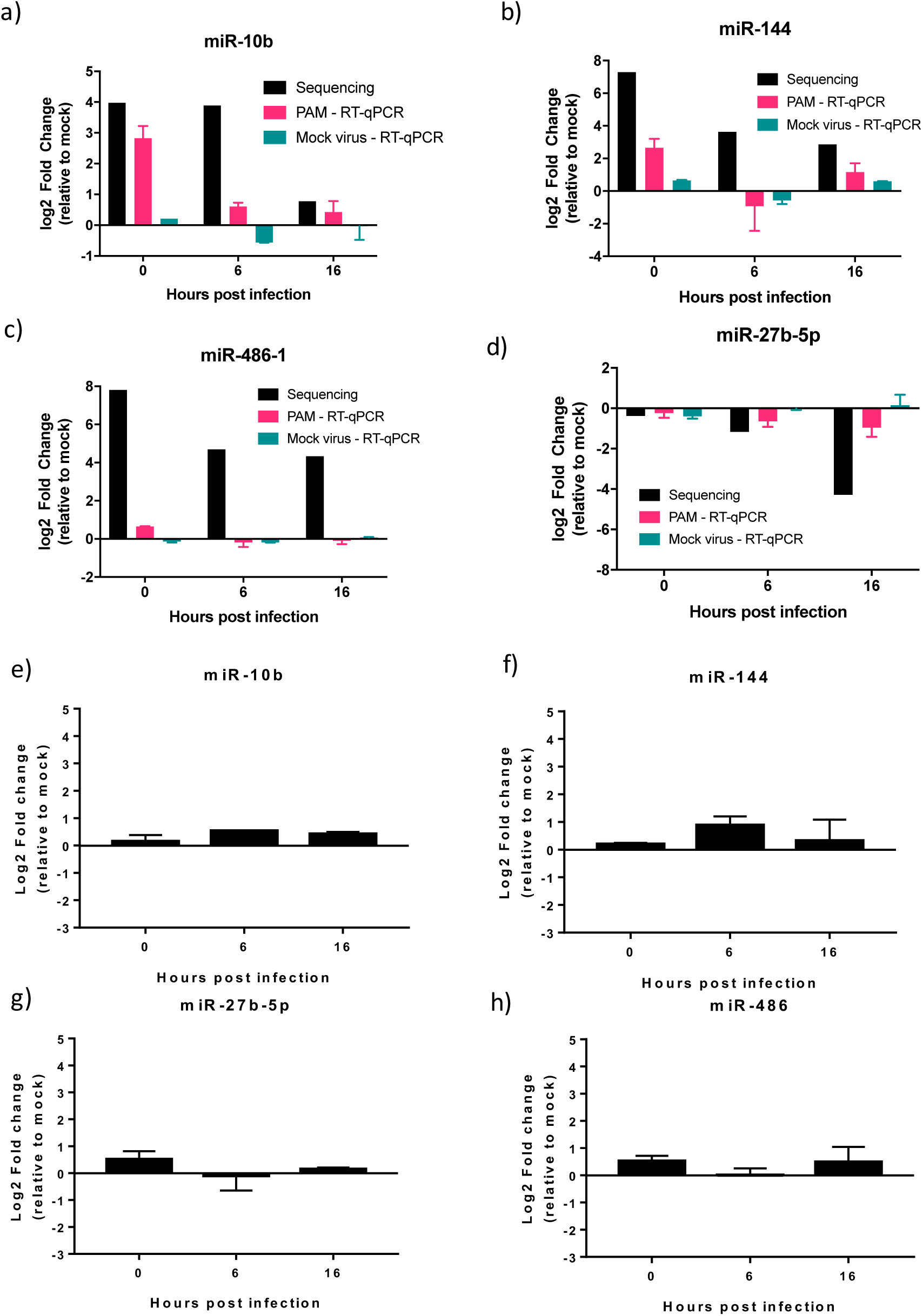
Differential expression of miR-10b and miR-144 in ASFV Benin 97/1 infected PAMs but not ASFV Ba71v infected Veros. **(a-d)** Changes in expression of miR-10b (a), miR-144 (b), miR-27b-5p (c) or miR-486-1 (d) in PAMs infected with ASFV Benin 97/1 or after the addition of a mock virus preparation at 0, 6 and 16 hpi were measured by RT-qPCR. Fold changes from miRNA sequencing are provided for comparison. (e-h) Changes in expression of miR-10b (e), miR-144 (f), miR-27b-5p (g) or miR-486-1 (h) in Vero cellss infected with ASFV Ba71v at 0, 6 and 16 hpi were measured by RT-qPCR. miRNA expression was normalised to U6 small RNA and fold change calculated by *Pfaffl* method. Data represents mean of 3 biological replicates, error bars indicate SEMs. Black bars: sequencing data results, pink bars: Benin 97/1 infected PAM RT-qPCR results, teal bars: Mock virus treated PAMs RT-qPCR results.

We also analysed miRNA abundance in response to ASFV infection in Vero cells using the Vero cell adapted stain, Ba71v. The expression pattern of the tested miRNAs did not follow that of Benin 97/1 infected PAMs with no substantial change in expression of any of the studied miRNAs (Fig 4e-f), with the log2 fold change not changing beyond ±1. Therefore, indicating either that the changes detected in PAMs with Benin 97/1 were cell-type specific or strain specific. Overall, we conclude that ASFV infection has a focused impact on the abundance of cellular miRNAs. This impact is limited to upregulation of a small number of miRNAs (miR-10b and miR-144) at very early time points during infection, specifically of Benin 97/1 infected PAMs.

### Identification of three ASFV encoded small non-coding RNAs

As ASFV does not mirror poxviruses by disrupting the host miRNA system on a widespread scale, there remains the potential for ASFV to utilise this system to encode its own miRNAs. To investigate this possibility, we aligned small RNA reads that did not map to known *Sus scrofa* sequences to the ASFV Benin 97/1 genome and the Ba71V genome, as this is the only isolate to have fully sequenced terminal inverted repeats. Plotting these aligned reads along the ASFV Benin 97/1 genome (Fig 5a) revealed a single peak of small RNA reads at approximately 82000 bp. This was also detected when aligned to Ba71v (Fig 5b). In addition, the Ba71v alignment revealed two peals of small RNA reads at 57bp and 170022 bp, indicating that these are located in genome termini. We termed these 3 small RNA sequences as ASFV small RNA 1, 2 and 3 (ASFVsRNA1, ASFVsRNA2, ASFVsRNA3). Both ASFVsRNA1 and ASFvsRNA3 were only detectable at 16 hpi with an average (mean) of approximately 100 reads per sample. ASFVsRNA2 was detectable at both 6 and 16 hpi with an average (mean) of 240 reads per sample. The sequences of the three small RNAs are shown in Fig 5c. Analysis of sequencing data from individual samples revealed that these small RNAs were only detected in infected samples (Fig 5 d, e, f) confirming that these sequences are virally derived. Due to the higher mean abundance of ASFVsRNA2 and its appearance at 6 hpi, this RNA was taken forward for further analysis.

**Figure 5:**
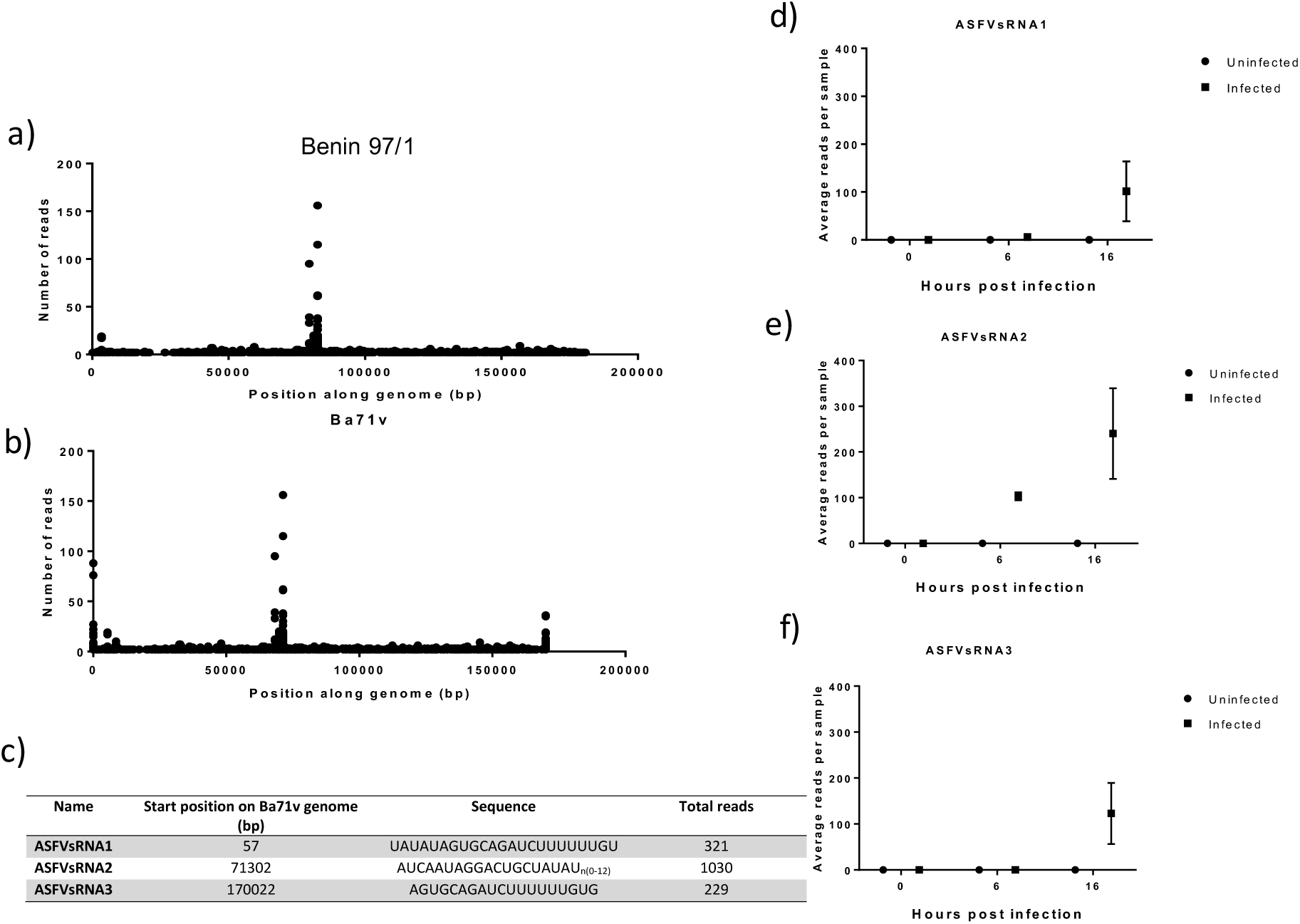
Mapping of unaligned small RNA reads to ASFV genome reveals peaks of small RNA sequences. a) Plot showing small RNA sequencing reads mapping to the ASFV Benin 97/1 genome, the position along the genome in bp is along the x-axis and the number of reads on the y-axis b) Plot of reads mapping to the Ba71v ASFV genome (this sequence includes the inverted terminal repeats). c) Table detailing the location and sequence of the peaks in reads. The average (mean) number of reads per sample at each time point is shown for d) ASFVsRNA1, e) ASFVsRNA2 and f) ASFVsRNA3. Error bars indicate SEMs.

### Characterisation of ASFV-encoded small RNA2

The sequence of ASFVsRNA2 aligns anti-sense (on the non-coding strand) to C147L (Fig 6a), the RNA polymerase subunit 6. This region of the gene is 100% conserved in all sequenced ASFV genomes (Fig 6b). Interestingly, ASFVsRNA2 had a variable number of 3’ U residues. A large proportion of reads contained 0 3’ uridines, 90% at 6 hpi, which decreased to 70% by 16 hpi (Fig 6c). The number of reads containing 1 – 12 3’ U residues increased over time from 10% at 6 hpi to 30% by 16 hpi (Fig 6c). The first 9 U residues are templated in the viral genome (depicted in Fig 6b) though around 10% of all reads have 10-12 at 16 hpi.

**Figure 6:**
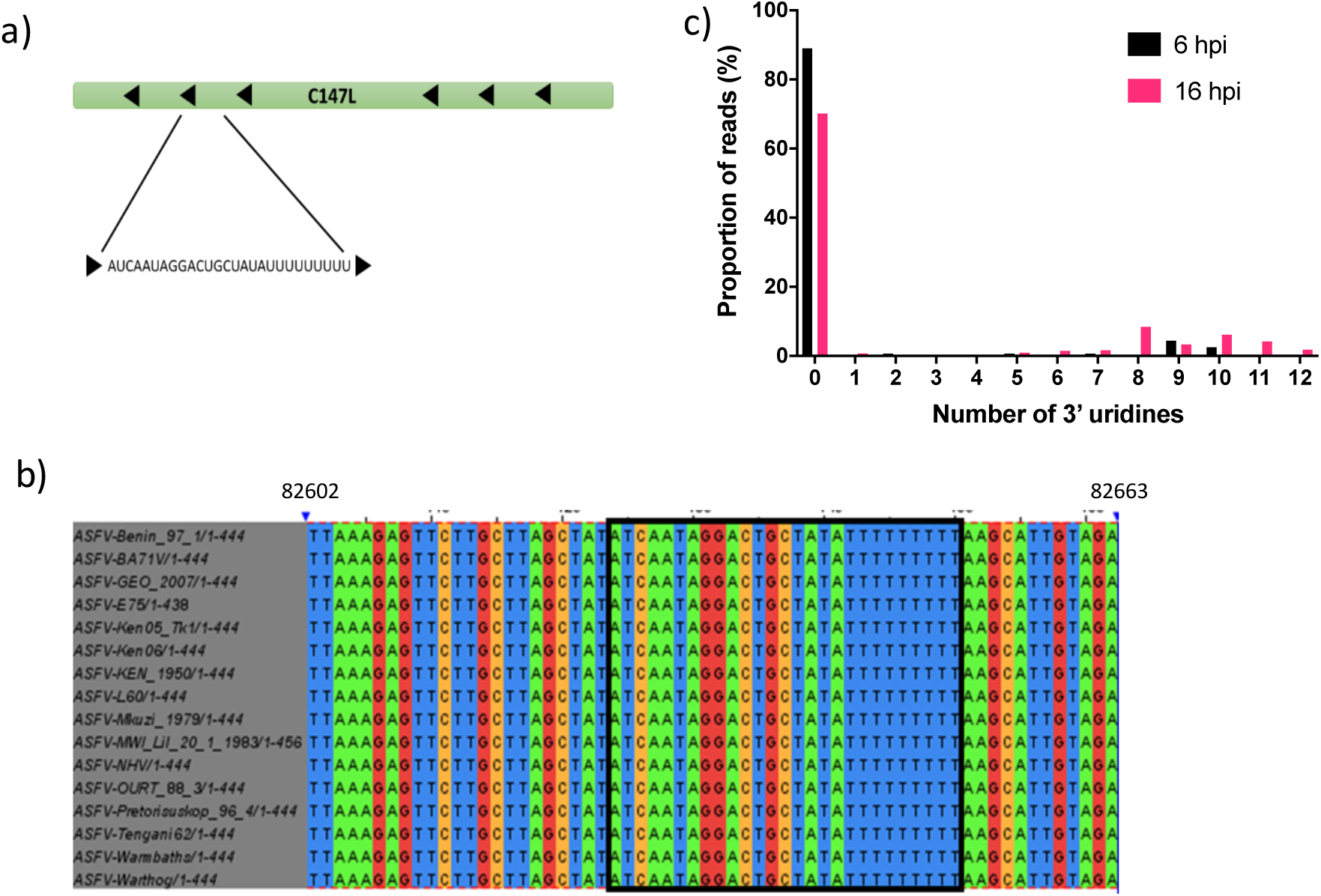
Alignment of ASFVsRNA2 to the ASFV genome. a) Location of the ASFVsRNA2 sequence in the C147L coding region, arrows represent coding direction. b) alignment of this region in multiple strains of ASFV, the boxed region highlights the ASFVsRNA2 sequence (alignment performed using clustal omega). Numbering is the genome position in ASFV Benin 97/1. c) The proportion of reads and the number of 3’ uridine residues of ASFVsRNA2.

Expression of the ASFVsRNA2 was validated using RT-qPCR. RNA from both ASFV Benin 97/1 infected PAMs and Ba71v infected VEROs was extracted at 0, 6 and 16 hpi. RT-qPCR was performed using a primer specific for ASFVsRNA2 (without the polyU sequence). ASFVsRNA2 was detectable at very low levels at 0 hpi (Fig 7a), with 40-Ct values 5 or below in both cell types. By 6 hpi, ASFVsRNA2 was readily detected with a mean 40-Ct value of at least 12 in both cell types, and even more abundant by 16 hpi, with the 40-Ct value increasing by at least 2. Next, in order to visualise ASFVsRNA2, northern blotting using a radiolabelled DNA probe perfectly complementary to ASFVsRNA2 was performed on the same cellular RNA samples. The 5S ribosomal RNA was used as a loading control. The probe detected a small RNA species in infected PAMs at 6 hpi, increasing in intensity by 16 hpi. A similar band also appeared in infected VEROs at 16 hpi (Fig 7b).

**Figure 7:**
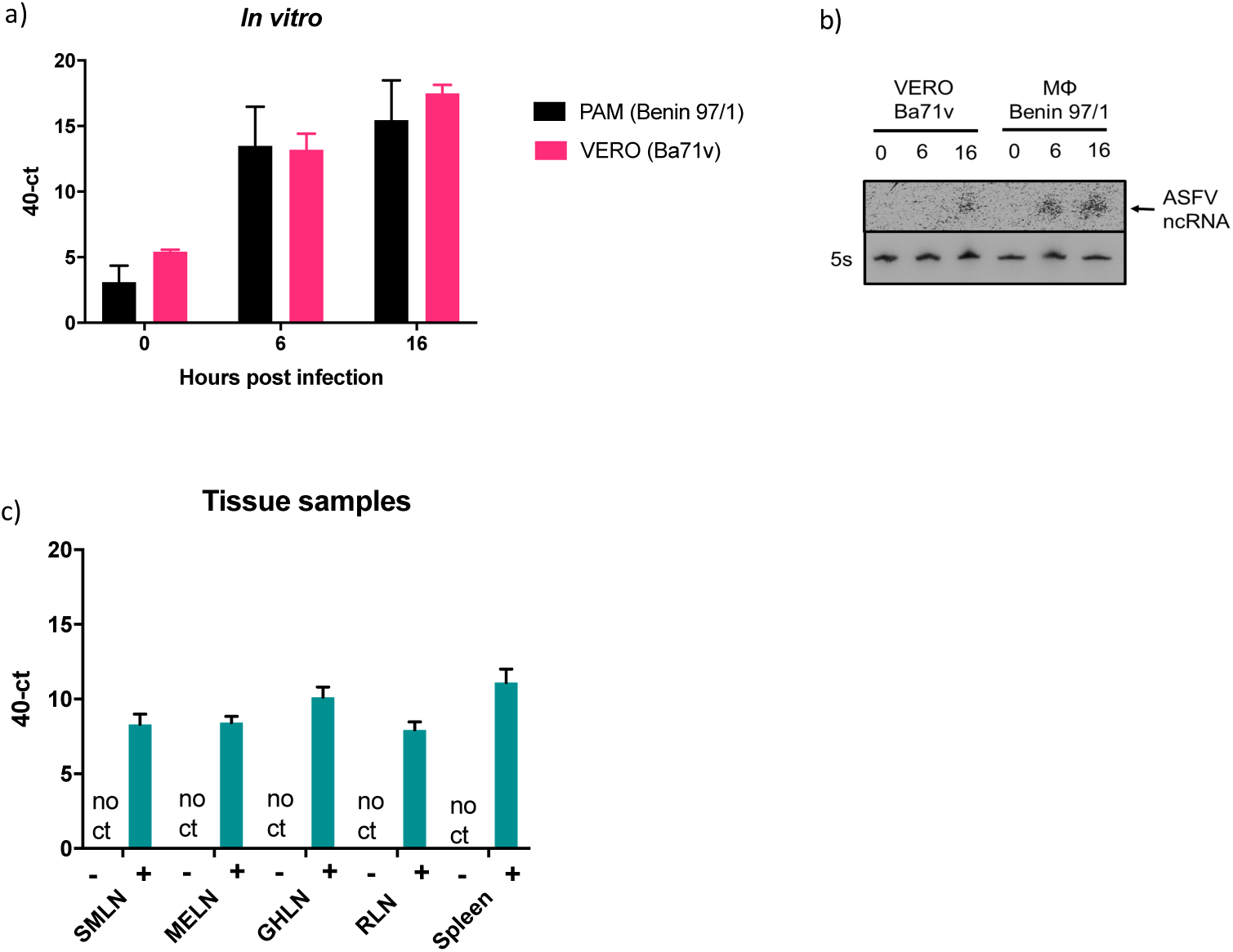
ASFVsRNA2 is expressed *in vivo*. Expression of ASFVsRNA2 was determined in ASFV-infected porcine macrophages and VERO cells at 0, 6 and 16 hpi by RT-qPCR (A) or Northern blot (B). 5s rRNA serves as a loading control. C) RT-qPCR analysis of ASFVsRNA2 expression in tissue sample from pigs infected with ASFV OURT88/1. (SMLN: submandibular lymph node, MELN: mesenteric lymph node, GHLN: gastrohepatic lymph node, RLN: renal lymph node). In panels a) and c), data represents mean of 3 biological replicates and error bars indicate SEMs.

To assess ASFVsRNA2 expression *in vivo*, RNA was analysed from tissues taken from pigs experimentally infected with ASFV. Outbred pigs were challenged with ASFV OURT88/1 and euthanised 5 days post challenge due to exhibiting moderate clinical signs consistent with ASF. At post-mortem examination the submandibular, mesenteric, gastrohepatic and renal lymph nodes as well as the spleen were taken from three pigs. Samples of the same tissues were also taken from one uninfected animal. RNA was extracted and RT-qPCR performed using the ASFVsRNA2 specific primer. ASFVsRNA2 was not detected in any of the samples taken from the uninfected animal. ASFVsRNA2 was detectable in all samples from animals infected with ASFV, with 40-Ct values ranging from 8 - 12 (Figure 7c), indicating that ASFVsRNA2 is produced during *in vivo* infection.

### ASFVsRNA2 is not produced through canonical miRNA biogenesis pathway

To investigate whether ASFVsRNA2 is a miRNA, we investigated whether it is produced through the canonical miRNA biogenesis pathway by assessing its loading into Argonaute 2 (Ago2), a key protein involved in miRNA biogenesis. Lysates were harvested from ASFV infected PAMs and incubated with either an anti-Ago2 antibody or non-immune rabbit sera as a negative control. Antibody-protein-RNA complexes were immunoprecipitated, and western blotting performed to identify enrichment of Ago2 in the anti-Ago2 antibody complexes compared to the non-immune antibody complexes and the non-precipitated cell lysate (Fig 8a). RNA was then extracted and RT-qPCR performed for ASFVsRNA2 and the miRNA miR-21, a highly expressed miRNA known to be loaded in RISC. Data was normalised to the small U6 RNA, which is not loaded in RISC. An eight log2 fold enrichment of miR-21 was detected in the complexes precipitated with the anti-AGO2 antibody, confirming the success of the IP (Fig 8b). However, ASFVsRNA2 was very poorly enriched (log2 fold-change of 1) after IP, indicating that ASFVsRNA2 is likely not produced through the canonical miRNA biogenesis pathway. We additionally used the miRNAFold program (22) to predict if the ASFVsRNA2 can form a miRNA stem-loop precursor. However, none of the *in silico* hairpin structures produced were convincing as miRNA hairpin precursors (data not shown).

**Figure 8:**
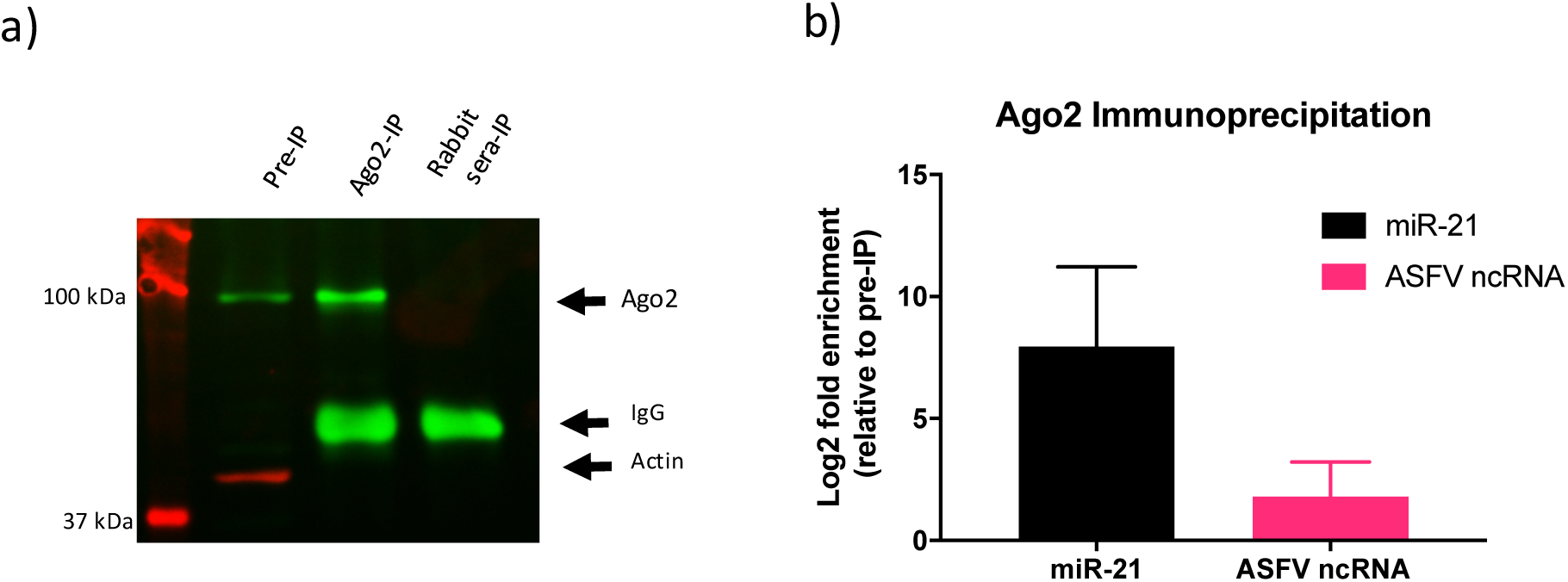
ASFVsRNA2 is not produced through the canonical miRNA biogenesis pathway. A) Western blot of ASFV-infected PAM lysates before and after Ago2 immunoprecipitation using anti Ago-2 and anti-actin antibodies. Pre-IP: lysate before immunoprecipitation, ago2-IP: Ago2 immunoprecipitation lysate. Rabbit-IP: pre-immune rabbit sera control immunoprecipitation b) The log2 fold enrichment after RT-qPCR for miR-21 and ASFVsRNA2 in the ago2 immunoprecipitation. Expression was normalised to U6 small RNA and fold change calculated by *Pfaffl* method. Data represents mean of 3 technical replicates and error bars indicate SEMs.

### Overexpression of ASFVsRNA2 reduces viral replication

We next sought to investigate whether ASFVsRNA2 performs a function during ASFV replication. We bypassed the canonical miRNA biogenesis pathway by synthesising single-stranded RNA mimics of the ASFVsRNA2 with and without the 3’ polyU sequence. These RNA mimics were stabilised with 2’-fluoro modifications as other sncRNAs have been shown to be functional and have targeting activity with this modification (23). The experiments were performed in Vero cells using Ba71v rather than PAMs due to the higher transfection efficiency of Vero cells. The ability of Vero cells to be both transfected with an RNA mimic and infected with ASFV was first examined. Cells were transfected with a Dy547-labelled miRNA mimic, miRDIAN and at 12 h later infected with ASFV Ba71v (After a further 24 h cells were fixed, permeabilized, and labelled with an antibody targeted to ASFV early CP204L/p30 protein. Analysis by confocal microscopy showed that all ASFV infected cells (p30 positive) were also transfected with the miRNA mimic, visible as red dots on the images (Fig 9a), indicating transfection and infection of the same cells had occurred.

**Figure 9:**
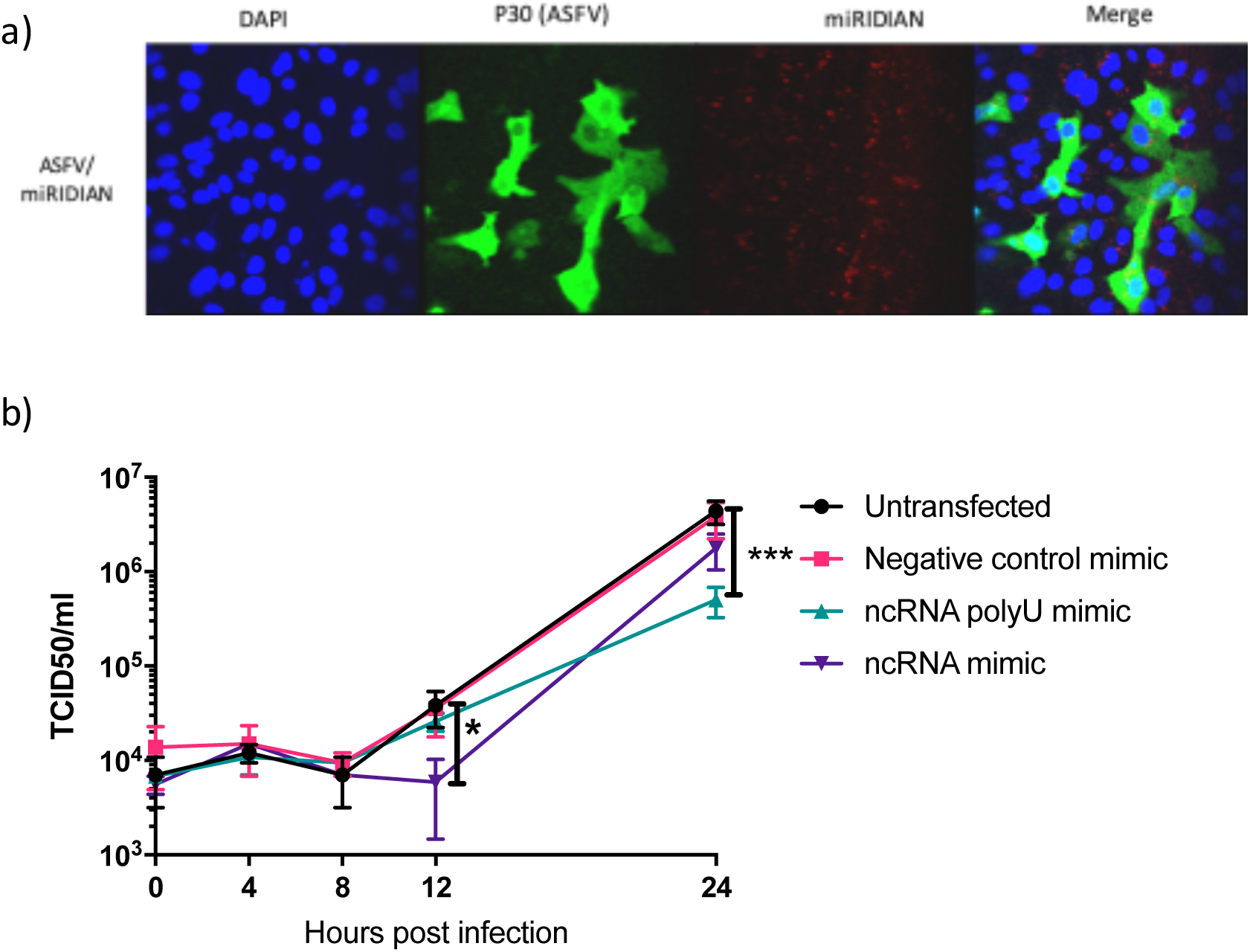
Overexpression of ASFVsRNA2leads to a reduction in ASFV replication. A) VERO cells transfected with miRIDIAN mimic transfection control with Dy547 for 24h were subsequently infected with ASFV Ba71v (MOI: 5). After 24h cells were fixed and labelled to confirm cells can be successfully transfected and infected. B) VERO cells were transfected with RNA mimics of ASFVsRNA2, with and without polyU sequence. After 24h cells were infected with ASFV Ba71v (MOI: 5) and supernatants were taken at 0, 4, 8, 12 and 24 hpi and titrated on Vero cells. TCID50 calculated by Spearman-Karber method. Error bars indicate SEMs *P= 0.0111, ***P=0.001, repeated measures two-way ANOVA. Representative of 2 biological repeats.

A single-step ASFV growth curve was then carried out on Vero cells transfected with either the ASFVsRNA2 mimic, ASFVsRNA2 polyU mimic, a negative control mimic or left untransfected. After 12 h, cells were infected with ASFV Ba71v and the amount of virus present determined at the time points shown by calculating the TCID50 (Fig 9b). The negative control mimic had no effect on viral replication, with no difference in viral TCID50 at any time point compared to the non-transfected cells. At 12 hpi, ASFV replication in cells transfected with the ASFVsRNA2 mimic had an approximately 0.5 log reduction in comparison to the non-transfected cells. Replication in these cells recovered by 24 hpi to a similar level as the negative control and non-transfected cells. Interestingly, the ASFVsRNA2 polyU mimic had a significant impact on replication only at 24 hpi, with an approximately 1 log reduction. The results therefore suggest that ASFV utilises both the 3’ uridylated and non-uridylated forms of this virally-encoded small RNA to regulate its own replication, and that the two forms are involved at different time points during replication.

## Discussion

With no effective vaccine or treatment, methods to control ASFV outbreaks are limited. As a result, after reaching the European Union (EU) in 2014 ASFV has subsequently spread throughout eastern Europe. In addition, the virus emerged in China in 2018 and has since spread rapidly across South East Asia with outbreaks declared in Vietnam, Cambodia, Mongolia and Hong Kong (24). ASFV in China has substantially impacted the food security in the world’s largest pig producer (25). Improving current ASFV vaccines and developing novel vaccines is therefore a priority. We therefore sought to investigate the interaction between ASFV and sncRNAs in order to gain more knowledge of ASFV host-pathogen interactions. Many viruses have been found to exploit and manipulate sncRNAs. This can range from subtle effects on specific host miRNAs to widespread disruption of miRNAs, such as in the case of poxviruses (17, 18). The mechanism of VACV-mediated miRNA polyadenylation is assumed to be mediated by the virally-encoded poly(A) polymerase (PAP), VP55 (17). The viral PAP is conserved throughout the NCLDV superfamily with the C475L gene identified as the putative ASFV PAP (26). We therefore first investigated whether ASFV shares the ability of poxviruses to induce miRNA polyadenylation and their subsequent decay. Both northern blotting and small RNA sequencing revealed no evidence of widespread miRNA polyadenylation and decay in ASFV-infected cells. This characteristic is therefore not conserved throughout the NCLDV superfamily and may be unique to poxviruses.

In contrast to poxviruses, this study found ASFV infection has only a modest impact on host miRNAs, with only 6 miRNAs identified by sequencing to be differentially expressed during ASFV infection, and only 3 of these robustly validated by RT-qPCR. This more subtle effect on miRNAs is more common in virus infections than the widespread disruption that poxviruses induce (27). For example, pseudorabies virus infected porcine dendritic cells led to the differential expression of only 8 miRNAs (28). A previous study examined miRNA expression in spleen and lymph node collected from pigs inoculated with a virulent and attenuated ASFV strain (19). This study identified differential regulation of 22 miRNAs in the spleen and 33 in the lymph node 3 days post inoculation. The only miRNA identified common to this study was miR-10b, which was found to be lightly downregulated between 3 and 7 dpi animals infected with a virulent ASFV strain. It is difficult to directly compare results as the previous study compared miRNA expression in pigs infected with a virulent strain to those infected with an attenuated strain, not uninfected animals. Additionally, our *in vitro* study focused on a single viral replication cycle over 16 hours whereas the *in vivo* experiment analysed viral infection over multiple days. However, the identification miR-10b as differentially expressed *in vivo* supports our theory that miR-10b plays a role during ASFV infection.

The upregulation of miR-10b occurred at 0 hpi, after the virus has been incubated on cells for only 1 h, but then rapidly decreased in abundance by 6 hpi. Further experiments using the porcine pestivirus classical swine fever virus (CSFV) also led to a similar pattern in miR-10b expression, with rapid upregulation of during the first hour of infection and subsequent decrease (data not shown). Both ASFV and CSFV are enveloped viruses and have been found to enter porcine macrophages via endocytosis (29) (30). In macrophages, miR-10b is known to target ATP binding cassette transporter A1 ABCA1 (31), which is involved in the regulation of cholesterol efflux. During infection with ASFV, cholesterol remodelling is essential in the establishment of productive viral infection and disruption of cellular cholesterol efflux leads to the impairment of virus entry and viral particles remained trapped in endosomes (32). This therefore suggests a link between miR-10b, ABCA1 and cholesterol efflux during endocytic-mediated entry of ASFV in macrophages.

A key finding of this study was identification and characterisation of an ASFV-encoded sncRNA, ASFVsRNA2. Due to their large genome size, DNA viruses have the coding capacity to encode miRNAs and many do, predominantly the nuclear replicating herpesviruses. As canonical miRNA biogenesis begins in the nucleus with host RNA Pol II transcription from the viral genome, it is assumed viral miRNA biogenesis requires a nuclear phase of viral replication. Indeed, virus-encoded miRNAs have been identified in NCLDVs that also have a nuclear phase of replication. These are in two Iridoviruses: Singapore grouper iridovirus (SGIV) (33) and Tiger Frog Virus (TGV) (34). Several studies have identified the presence of ASFV genomes in the nucleus at early time points in infection (35), this nuclear phase remains poorly understood but it indicates the possibility for ASFV to encode miRNAs.

Our study identified ASFVsRNA2 in both PAMs and tissues from pigs with ASFV (Fig 7a-c). A previous *in vivo* study concluded that ASFV does not express viral miRNAs in experimentally infected pigs (21). However, the previous study restricted its analysis to miRNA only based on predictions of pre-cursor miRNA structures and did not consider other classes of sncRNA. Since a number of virus families encode different classes of sncRNAs (reviewed in (14)) we chose to sequence the small RNA fraction from ASFV-infected cells without bias, and therefore allowed us to report the first identification of an ASFV-encoded sncRNA.

Our research has indicated that ASFVsRNA2 does not fit the classic miRNA biogenesis pathway as it does not have an identifiable hairpin precursor and fails to enrich in an Ago2 immunoprecipitate. However, it is known that a number of viral miRNAs are produced though non-canonical pathways. (36). For example, miRNAs encoded by murine γ-herpesvirus 68 (MHV68) in which the pre-cursor miRNAs are located in a tRNA-like structure. (37). Despite this variety of biogenesis pathways, miRNAs are required to be loaded into an Ago protein in order to function. Pigs, like other mammals, have been found to encode 4 Ago proteins (Ago1-4) (38), though only Ago2 is catalytically active and has the ability to cleave target mRNA (39). It has been reported that miRNAs are not sorted into distinct human Ago proteins (40) and so miRNAs would be expected to be found in all Ago proteins. Therefore, our inability to detect ASFVsRNA2 enrichment in an Ago2 immunoprecipitation indicates that ASFVsRNA2 is not a miRNA.

The presence of polyuridine (polyU) residues on the 3’ end of ASFVsRNA2 is intriguing. The first 9 of these uridine residues are templated in the ASFV genome, though a small percentage of reads at 16 hpi have 10-12 uridine residues. It is likely that the variation in the number of templated uridine residues is a result of ASFVsRNA2 biogenesis. ASFV transcription termination takes place at a conserved motif of seven or more consecutive thymidylate residues (41) and this poly(T) motif is retained in the mRNA (42). As the location ASFVsRNA2 maps to in the genome does not correspond to a 3’ end of any known gene, this polyU motif could be the termination signal for ASFVsRNA2 transcription. The non-templated additions may indicate a modification to ASFVsRNA2 as the main function of non-coding RNA uridylation is to promote it degradation (43). However, it can also have a number of other functions. For example, uridylation of the miRNA let-7 pre-cursor is required for let-7 biogenesis (44). It can also control the activity of non-coding RNA, for instance uridylation of miR-26 prevents it binding to its target IL-6 transcript but does not affect miRNA stability (45). The functional significance of the uridine residues, templated or not, on the 3’ end of ASFvsRNA2 is currently unknown, though our work has indicated it is of importance.

Overexpression of ASFVsRNA2 reduced ASFV replication in Vero cells, suggesting it has a role in the control of viral replication. This relatively modest effect, between 0.5-1 log reduction, is consistent with sncRNA gene targeting since miRNAs usually regulate gene transcript levels by less than 2-fold (46). Even though we do not believe ASFVsRNA2 to be a miRNA, the predominant function of all classes of sncRNAs is to induce gene silencing (47). Therefore, we would also expect ASFVsRNA2 to have only a modest effect. Many viruses are known to encode activators and repressors to finely control their life cycle in order to replicate in the face of the host immune response. An interesting example of this is the Human Cytomegalovirus (HCMV) encoded miRNA, miR-UL112-1, which has been found to downregulate multiple viral genes involved in replication, leading to a decrease in genomic viral DNA (48).

In summary, we have found that ASFV infection has a very modest impact on host sncRNA and instead utilises this system to encode its own, ASFVsRNA2, which retards viral replication. The discovery of an ASFV-encoded sncRNA has added another level to the knowledge of ASFV replication and opens up the possibility of sncRNA-based mechanisms to develop the next generation of ASFV vaccines.

## Materials and methods

### Cells and viruses

African green monkey epithelial cells (Vero) were grown in Dulbecco’s modified Eagle’s medium (DMEM) (Life Technologies), containing 100 units/ml penicillin, 100μg/ml streptomycin (pen/strep) (Sigma), and 10% foetal bovine serum (FBS) (Life Technologies). Cells were maintained at 37°C with 5% CO_2_ and were passaged regularly to maintain viability. Primary porcine cells were derived from 4 week old Large White outbred pigs. Porcine alveolar macrophages (PAMs) were obtained by lung lavage with PBS and maintained in RPMI 1640 GlutaMAX™ (Life Technologies) containing 100 units/ml penicillin, 100μg/ml streptomycin (pen/strep) (Sigma) and 10% porcine serum (BioSera). Bone marrow cells were prepared from femur bones and maintained in Earle’s balanced salt solution (EBSS) 100 units/ml penicillin, 100μg/ml streptomycin (pen/strep) (Sigma), 10% porcine serum (BioSera) and 10mM HEPES. The pathogenic ASFV Benin 97/1, previously described in (49) was grown in primary porcine bone marrow cells and the tissue-culture adapted ASFV BA71V, previously described in (50) was grown in Vero cells. Experiments involving VACV used the Western Reserve (VACV-WR) strain, prepared by purification on 36% (wt/vol) sucrose cushion. Experiments involving CSFV used the Brescia strain, kindly provided by Dr Julian Seago (The Pirbright institute)

### Animal experiments and Ethics Statement

Animal experiments were carried out under the Home Office Animals (Scientific Procedures) Act (1986) (ASPA) and were approved by the Animal Welfare and Ethical Review Board (AWERB) of The Pirbright Institute. The animals were housed in accordance with the Code of Practice for the Housing and Care of Animals Bred, Supplied or Used for Scientific Purposes, and bedding and species-specific enrichment were provided throughout the study to ensure high standards of welfare. Through careful monitoring, pigs that reached the scientific or humane endpoints of the studies were euthanised by an overdose of anaesthetic. All procedures were conducted by Personal Licence holders who were trained and competent and under the auspices of Project Licences. Female Landrace × Large white (Yorkshire) × Hampshire pigs were obtained from a high health farm in the UK. Animals were challenged intramuscularly in the rump with 10,000 HAD of the OUR T88/1 strain of ASFV. Tissue samples were collected from three pigs at post mortem five days post challenge.

### Preparation of RNA samples

Confluent 6-well plates of Veros and PAMs were infected or mock-infected with ASFV or VACV at an MOI of 10 for 1h at 37°C. The inoculum was removed (0h time point), cells were washed 3x in PBS and media, containing 2.5% serum, was replaced. At 0, 6 and 24 hpi cells were harvested into an appropriate volume of QIAzol lysis reagent (Qiagen) and RNA prepared using the miRNeasy Mini Kit (Qiagen). For RNA extraction from animal tissues, the tissues were harvested into RNAlater (Life Technologies) and stored at −80°C. 50mg of tissue was added to 700μl Qiazol lysis reagent (Qiagen) and homogenised using tissue grinding lysate matrix beads (MP Biomedicals). RNA was prepared using the miRNeasy Mini Kit (Qiagen), including an on-column DNase digest (Qiagen).

### Northern blot analysis

Northern blotting was carried as described in (18). Briefly, 5µg RNA was mixed with 2x TBE-UREA buffer (Novex) and heated at 70°C for 3 min. Samples were run on a 15% polyacrylamide TBE-UREA gel (BioRad). The gel was then transferred to a solution of 0.5 x TBE containing 10,000 x SYBR gold (Invitrogen) and visualised under UV light to check for equal loading. The RNA was transferred to a Hybond N^+^ membrane (Amersham) using a semi-dry transfer machine and cross-linked as described in (51). DNA probes perfectly complementary to a small RNA were prepared by labelling with P^32^ using the mirVana probe and marker kit (Ambion) Membranes were pre-hybridised in ULTRAhyb hybridisation buffer (Ambion) for 1hr at 42°C prior to incubation with a P^32^-labelled DNA probe, overnight at 42°C. Membranes were then washed twice in 2x SSC with 0.1% SDS for 15 min and laid against a phosphorimage screen for 4-6hrs. Labelling was detected using a Typhoon FLA 7000 phosphorimager (GE Healthcare). To strip the membrane for re-probing it was washed in a solution of boiling 0.1% SDS for 30min.

### Small RNA sequencing

Integrity of the RNA was measured on a Bioanalyzer (Agilent), with all samples having a RIN value of 8 or above. Small RNA sequencing libraries were prepared using CleanTag small RNA kit (TriLink). Libraries were pooled, gel purified on a 5% polyacrylamide TBE gel (BioRad) and sequenced on an Illumina HiSeq (Edinburgh Genomics). An average of 7013808 ± 3522153 reads per samples were generated. Quality of reads was assessed using FASTQC and adapters trimmed using cutadapt software. Sequences were collapsed within each sample to generate a non-redundant set of fasta sequences. Singletons were not included. The reference used for alignment was version 10.2 of the *Sus scrofa* genome obtained from Ensembl, only full-length perfect match (FLPM) sequences were counted. Sequences aligning to the genome were subsequently used as input for a mirDeep2 analysis. Alignments were performed using a non-current version of bowtie-based Perl script (mapper.pl) that forms part of the mirDeep2 software package. The mirDeep2 version was 2.0.0.4 and the bowtie version was 0.12.5. Parameters used were *-o 20 -l 17 -r 100 –c*. The analysis used *Sus scrofa* mature (3p and 5p forms) and precursor sequences obtained from mirBase (release 21). Small RNA reads that did not map to *Sus scrofa* sequences on mirBase were aligned to the ASFV BA71V genome (Genbank: U18466.2) and ASFV Benin 97/1 genome (Genbank: AM712239.1). Sequencing data was deposited in NCBI GEO database under accession GSE115512

### Differential expression analysis

Initial raw counts were filtered to only include those with an average of 5 reads or more. The counts within each sample were converted to abundances, which were (1) multiplied by one million to generate a reads set, (2) one count added to all to preclude zero counts instances, and (3) the resultant values converted to log2 and quantile normalised. Pairwise comparisons of sample groups were performed on the normalised tag counts using linear modeling (Bioconductor *limma* package). A series of 6 group-wise comparisons using empirical Bayesian approaches was undertaken to identify differences (fold changes). Significance values are controlled for false discovery, yielding a more rigorous adjusted P value.

### Quantitative reverse transcription PCR (RT-qPCR)

cDNA was generated using the miScript II RT Kit (Qiagen). qPCR was performed using the miScript SYBR Green PCR Kit (Qiagen) using miScript Primer Assays for miRNAs and the using the QuantiTect SYBR Green PCR kit (Qiagen) for mRNA in a Stratagene Mx3005P qPCR machine (Agilent). All qPCRs were performed in duplicate. Hs_RNU6-2_11 miScript Primer was used as a reference gene for sncRNA data normalisation and 18s rRNA was used for mRNA data normalisation. The PCR efficiency of each miScript Primer was determined by standard curve and the log2 fold change calculate by the *Pfaffl* method.

### Immunoprecipitation and Western blotting

Lysates were prepared by washing cells x2 in ice-cold PBS before addition of an appropriate volume of RIPA lysis buffer (ThermoFisher) supplemented with protease inhibitor (Complete™ Protease Inhibitor Cocktail Tablets, Roche) and for immunoprecipitation, RNase inhibitor (ThermoFisher). Protein concentrations were determined using BCA Protein assay Kit (Thermo Scientific Pierce). For immunoprecipitation, equal amounts of protein were incubated with either a rabbit polyclonal anti-Ago2 antibody or pre-immune rabbit sera overnight at 4°C. This was followed by a 1h incubation at room temperature with 25μl Pierce protein A Magnetic Beads (Thermo Scientific). The beads were washed and resuspended then either prepared for Western blotting or added to Qiazol lysis reagent for RNA extraction. For western blotting, lysates or bead suspensions were prepared by mixing 20µg of protein with 2X Protein Sample Loading Buffer (Li-Cor) and heated to 98°C for 5min. Samples were loaded onto a 15% polyacrylamide resolving gel, layered with a 5% stacking gel, alongside a pre-stained protein ladder (Biorad) and the proteins separated by electrophoresis. Proteins were transferred to a PVDF membrane using a wet transfer technique. Membranes were first blocked for 1 hr at room temperature in a 1:1 mixture of PBS and Odyssey blocking buffer (Li-Cor) then incubated with the primary antibodies diluted in Odyssey blocking buffer and 0.1% Tween at 4°C overnight. Membranes were washed 4 x for 5min in PBS containing 0.1% Tween before being incubated with the secondary antibodies diluted in Odyssey blocking buffer and 0.1% Tween for 45min at room temperature. Primary antibodies used were rabbit anti-Ago2 (Kindly provided by F. Grey), Rat anti-HA, rabbit anti-actin (Cell signalling) and mouse anti-actin (Cell signalling). The secondary antibodies were DyLight 680 and 800 (Cell Signalling) Membranes underwent a further 4 5min washes in PBS-T and were then visualised on a G:Box (Syngene).

### Plasmid and RNA mimic transfection

Vero cells were seeded at an appropriate density 24 h before transfection. Both plasmid DNA and RNA mimics were transfected using Transit-X2 (Mirus), following manufacturer’s protocol. In brief 500ng DNA and/or 10μM RNA mimic was diluted in 50μl optiMEM (LifeTech), 1.5μl transfection reagent was added and incubated for 15 min at room temperature before adding to 1 well of a 24-well plate. The single-stranded RNA mimics were synthesised (Sigma) with 5’-phosphorylation and 2’-fluoro modification for stability. The sequences of the RNA mimics were as follows, ASFVsRNA2 mimic: AUCAAUAGGACUGCUAUA, ASFVsRNA2 polyU mimic: AUCAAUAGGACUGCUAUAUUUUUUU and negative control: UUCUCCGAACGUGUCACGU. The miRIDIAN microRNA mimic transfection control with Dy547 was sourced from Dharmacon. Mimics were transfected to give a final concentration of 25nM per well.

### One-step growth curve

Vero cells were transfected with RNA mimics as described above and incubated for 14 h at 37°C. Cells were then infected with ASFV Ba71v at an MOI of 5 for 1 hr at 37°C, washed x3 and media replaced. Virus was harvested at 0, 4, 8, 12 and 24 hpi by scraping the cells into the media and freeze-thawing x3. Viral titres were determined by immunofluorescence TCID50 assay on Vero cells using an antibody against ASFV P30 protein. TCID50 was calculated using the Spearman-Karber method.

### Immunofluorescence

Cells were fixed after 3x wash in ice-cold PBS with 10% formalin and incubated for 30min at room temperature. Cells were washed again 3x in PBS and permeabilised by the addition of 0.2% triton-X100 diluted in PBS for 5min at room temperature and washed a further 3x in PBS. The primary antibody (mouse anti-ASFV P30) was diluted in PBS with 2% FBS and incubated with the cells in a humidity chamber for 1hr. Cells were then washed 3x in PBS with 2% FBS and incubated with secondary antibody (Alexa Fluor 488-conjugated goat anti-mouse IgG) and fluorescently-tagged phalloidin (Molecular Probes), in a humidity chamber for 1hr. Coverslips were stained with 300nM DAPI (Life Technologies) for 5 min and then rinsed 3x in PBS with a final rinse in dH_2_O before being mounted onto a microscope slide, using Vectashield (Vector Labs).

## Data availability

Raw data available on NCBI GEO database under accession GSE115512

